# Telomere length and oxidative stress variations in a murine model of Alzheimer’s disease progression

**DOI:** 10.1101/684365

**Authors:** Katia Martínez-González, Azul Islas-Hernández, José Darío Martínez-Ezquerro, Federico Bermúdez-Rattoni, Paola Garcia-delaTorre

**Author notes:** Corresponding author: Paola Garcia-delaTorre, Cuauhtémoc 330, Col. Doctores, Delegación Cuauhtémoc, Ciudad de México, México.

## Abstract

Alzheimer’s Disease (AD) is the most common cause of dementia and aging is its major risk factor. Changes in telomere length have been associated with aging and some degenerative diseases. Our aim was to explore some of the molecular changes caused by the progression of AD in a transgenic murine model (3xTg-AD; B6; 129-Psen1 <tm1Mpm> Tg (APPSwe, tauP301L) 1Lfa). Telomere length was assessed by qPCR in both brain tissue and peripheral blood cells and compared between three age groups: 5, 9, and 13 months. In addition, a possible effect of oxidative stress on telomere length and AD progression was explored. Shorter telomeres were found in blood cells of older transgenic mice compared to younger and wild type mice but no changes in telomere length in the hippocampus. An increase in oxidative stress with age was found for all strains but no correlation was found between oxidative stress and shorter telomere length for transgenic mice. Telomere length and oxidative stress are affected by AD progression in the 3xTg-AD murine model. Changes in blood cells are more noticeable than changes in brain tissue, suggesting that systemic changes can be detected early in the disease in this murine model.

## Introduction

Alzheimer’s disease (AD), the most common cause of dementia, is defined by progressive and irreversible neurodegeneration of the central nervous system that eventually leads to the gradual decline of cognitive function (Czech *et al.*, 2000; Oddo *et al.*, 2003). The main feature of this disease is the deposition of amyloid-β (Aβ) protein in the extracellular space of the cerebral cortex and the walls of cerebral blood vessels. It has traditionally been thought that the Aβ deposited in the brain originates from the brain itself. However, it has recently been speculated that it may also come from Aβ at the periphery circulating in the blood. It is known that peripheral exposure to Aβ-rich brain extracts can also induce cerebral Aβ deposits in transgenic mice (Meyer-Luehmann *et al.*, 2006). In this regard, insoluble Aβ oligomers have been experimentally shown to cause memory dysfunction, inhibit LTP, and prolong LTD (Ding *et al.*, 2019).

In addition to these neuronal lesions, some biomarkers of genomic instability have been found in patients with AD, including micronuclei (markers of loss and breakage of chromosomes), aneuploidies of chromosome 21, shortening of telomeres in lymphocytes and fibroblasts (Thomas *et al.*, 2008), and increased oxidative stress markers (Butterfield & Sultana, 2011).

Telomeres are essential non-coding deoxyribonucleic acid (DNA)-protein complexes that cap the ends of the linear chromosomal DNA protecting the genome from damage. In mitotic cells, telomeres can shorten with each division, unless this can be counteracted or reversed by the telomere-lengthening enzyme, telomerase (Blackburn, 1991; Wolkowitz *et al.*, 2011), telomere length regulation in mammalian cells is complex, telomere length differs between tissues (Prowse & Greider, 1995) or even cells (Friedrich *et al.*, 2000).

An increase in tissue cell proliferation results in the progressive shortening of telomeres, which triggers a DNA damage response that if repaired, restores proliferation. On the contrary, if the deterioration is irreversible, the cell keeps the cell cycle at halt and can be directed to apoptosis or cellular senescence (Meeker *et al.*, 2004; Campisi, 2011). Telomere length depends on several factors such as the speed of degradation and the speed and time of action of telomerase in each chromosome (Cawthon *et al.*, 2003; Jaskelioff *et al.*, 2011). Hence, telomere length decreases differentially with time in all mitotic tissues.

The brains of patients diagnosed with AD show a significant extent of oxidative damage associated with the abnormal marked accumulation of β-amyloid and the deposition of neurofibrillary tangles (Christen, 2000); stress contributes to a significant decrease in telomerase activity and consequently causes an increase in the telomere-shortening rate (Kurz *et al.*, 2004). Biochemical studies show TTAGGG repeats are preferred sites for iron binding and iron mediated Fenton reactions, which generate hydroxyl radicals that induce the 5’ cleavage of GGG (Oikawa *et al.*, 2001). There is a minimal shortening of less than 20 bp per cell division in cells with high antioxidative capacity and this rate increases in cells with lower antioxidative defense. Cultivating cells under enhanced oxidative stress like mild hyperoxia (40% normobaric oxygen) shortens the telomeres prematurely and reduces the replicative lifespan accordingly (Saretzki & Von Zglinicki, 2002).

Telomere length has been associated with life span since it reflects the number of times a cell has divided (Boonekamp *et al.*, 2013); during aging, 50-150bp of telomeric DNA is lost with each proliferation cycle (Hochstrasser *et al.*, 2012). It also has been associated with disease conditions such as mental stress, obesity, smoking, type 2 diabetes mellitus, ischemic heart diseases, AD, and Parkinson’s disease (Harley *et al.*, 1990; Allsopp *et al.*, 1992; Zakian, 1995; Brouilette *et al.*, 2003).

Specifically, it has been reported that patients with AD show shorter telomeres in peripheral blood mononuclear cells, monocytes, T and B cells (Panossian *et al.*, 2003), buccal cells, and leukocytes compared to controls (Thomas *et al.*, 2008). Also, significant differences in leukocyte telomere length (LTL) between controls, amnestic mild cognitive impairment, and AD patients has been recently reported (Scarabino *et al.*, 2017). However, there are no reports, to our knowledge, of how and if progression of AD affects telomere length. Hence, we decided to evaluate changes in telomere length and oxidative stress due to the progression of AD in a murine model. Since our model has a genetic disposition to develop AD we expected to see an intrinsic effect of disease progression (changes in oxidative stress) on telomere length.

## Materials and methods

### Ethical standards

All procedures were performed in accordance with the current rulings in Mexican law (NOM-062-ZOO-1999) and with the approval of the local Science and Ethics Committees (R-2012-785-049).

### Animals

Homozygous 3xTg-AD (3xTg-AD; B6; 129-Psen1 <tm1Mpm> Tg (APPSwe, tauP301L) 1Lfa) male mice were used as an AD murine model, and B6129SF2/J WT male mice were used as controls since this is the genetic background originally described for the transgenic mice used. Mice were purchased from Jackson Laboratories and housed in the *Bioterio de la Coordinación de Investigación en Salud*. Mice were housed in a 12:12 light/dark cycle at 20-22°C with water and food *ad libitum*. All animals were genotyped. Animals of three different ages were used for this project 5, 9, and 13 months. Groups for telomere length measurement were as follows: 5mo 3xTgAD n=7, 5mo WT n=12; 9mo 3xTgAD n=9, 9mo WT n=12; 13mo 3xTgAD n=8, and 13mo WT n=12. Groups for oxidative stress measurements were as follows: 5mo 3xTgAD n= 6, 5mo WT n=6; 9mo 3xTgAD n= 6, and 9mo WT n= 6.

### Biological samples and DNA extraction

Blood samples were obtained from the mandibular vein (~0.2ml) and treated with an erythrocyte lysis buffer. After the last blood sample was obtained from each group of age, mice were sacrificed with an overdose of pentobarbital administered intraperitoneally. The animals were then decapitated with a guillotine, the brain was carefully removed, tissue from the hippocampus was dissected and homogenized with lysis buffer. DNA was extracted according to the manufacturer’s instructions (Thermo Scientific DNA extraction kit, K0512). Purified DNA samples were stored at −70 °C until use.

### Telomere length assessment

All measurements were performed on samples from 5, 9, and 13 month-old mice from both strains. We followed the qPCR (StepOnePlus Real-Time PCR System, Applied Biosystems) method published by O’Callaghan and Fenech (O’Callaghan & Fenech, 2011), an absolute quantification method that introduced an oligomer standard. A standard curve was used to determine telomere length; the 36B4 housekeeping gene was used as an endogenous calibrator. The number of copies of telomeric repeats was determined by the standard curve of telomere standard, while the standard curve of 36B4 STD was used as control, oligomers was HPLC-purified synthetized.

The Ct values of each sample were extrapolated in their corresponding curves by a linear regression test. The Maxima SYBR Green/ROX qPCR Master Mix 2X (ThermoScientific, California, USA) was used. The cycling conditions for both genes were: 10 minutes at 95°C, followed by 40 cycles of 95°C for 15 seconds, 60°C for 1 minute, followed by a melting curve. Results are expressed as absolute telomere length (aTL); Kb telomere/diploid genome and were obtained as suggested by O’Callaghan and Fenech (O’Callaghan & Fenech, 2011).

### Oxidative Stress measurements

All measurements were performed on samples from 5 and 9 month-old mice from both strains. No 13 month-old mice were used for these measurements.

### Reactive oxygen species (ROS)

ROS were measured by the oxidation of 2-7-dichlorofluorescein (DCFH) to the fluorescent oxidized compound 2-7-dichlorofluorescein (DCF) by the presence of hydrogen peroxide. Several reactive intermediates can oxidize DCFH, so it cannot be used to determine the presence of a specific reactive species.

To evaluate the formation of reactive oxygen species (ROS) by fluorometry, 60μl of homogenates of the hippocampus and whole blood cells were used. A final volume of 200μl was obtained with 1x PBS buffer, 10μL of a DCFDH-diacetate at 75 μM. Samples were incubated in the dark for 30 min at 37°C, then centrifuged at 10,000 rpm for 10 min; the supernatants were read in a fluorometer at an excitation wavelength of 480 nm and emission of 532 nm. Results are expressed in nM of DCFH in mg of tissue samples or nM of DCFH in μL of serum.

### Lipid peroxidation

Lipid peroxidation was measured by the production of malondialdehyde (MDA); 120μL of the homogenate of the hippocampus or 60μL whole blood cells were mixed with 60μL of 1x PBS and 120μL of the TBA reagent (0.375g of TBA + 15g of trichloroacetic acid + 2.54ml of concentrated HCl). Samples were placed in a boiling bath (94 °C) for 20 min subsequently centrifuged at 10,500 rpm for 15 min. The optical density of the supernatant was determined (Biotek plate reader) at a wavelength of 532 nm. Results are expressed as μM of MDA in μL of serum and μM of MDA in mg of tissue sample.

### Mitochondrial viability

Mitochondrial viability was measured by the reduction of 3-(4,5-dimethylthiazol-2-yl)bromide-2,5-diphenyltetrazole (MTT nM/μL); 100μL of brain tissue homogenate was used, to which 10μL of an MTT solution (5mg / mL) was added, and incubated for 30 minutes at 37 °C in the dark. After incubation, samples were centrifuged at 12,000 rpm for 3 min. The supernatant was removed, and 500 μL of isopropanol-acid was added to the pellet where it was suspended. Samples were read at a wavelength of 560 nm in a plate reader. Results are expressed as μM of MTT in mg of tissue.

### Statistical analysis

The data analysis was carried out in Prism 6.0e and SPSS 21 commercial software. As a first approach to explore the molecular changes in the progression of Alzheimer’s disease in a murine transgenic model (3xTgAD), we measured absolute telomere length (aTL; kb) at 5, 9, and 13 months, in addition to reactive oxygen species (DCFH; nM/μL), lipid peroxidation (MDA; μM/μL), and mitochondrial functionality (MTT; nM/μL) at 5 and 9 months. All measurements were carried out on the hippocampus homogenate and whole blood cells. Quantitative variables are presented as the arithmetic median and maximums and minimums per age group: 3, 5, and 13 months.

Scattergrams show the distributions of aTL among age groups, p-values were calculated by Kruskal-Wallis non-parametric test, and when statistical significance between groups was obtained, we performed a post-hoc test (Dunn’s test) to discriminate the specific groups with significantly different medians that differ from the others.

Differences between ROS, MDA, and MTT among age groups are represented as scattergrams. A Kruskal-Wallis (KW) non-parametric test was used to calculate p-values when statistical significance between groups was obtained, we performed a post-hoc test (Dunn’s test) to discriminate the specific groups with significantly different medians that differ from the others; in some cases, a Mann-Whitney test was used to compare between two groups.

To determine the possible relationship and strength of the relationship between ROS, MDA, and MTT with absolute telomere length among whole blood cells or hippocampus homogenate from either AD (3xTgAD) or Wt (B6129SF2/J) mice at 5 or 9 months, we performed a Spearman’s rank-order correlation (rs) analysis and considered the statistical significance of the correlation according to p-values less than 0.05.

### Availability of supporting data

https://osf.io/pygce/?view_only=06b96ac7527a4385b765bf3b5036d77b

### Competing interests

We declare no conflict of interest.

## Results

We measured telomere length and oxidative stress in the progression of AD in a murine model. Table 1 shows the mean values of telomere length, measured in whole blood cells and hippocampus at 5, 9, and 13 months, as well as ROS, lipid peroxidation, and mitochondrial viability measured at 5 and 9 months, for both transgenic and wild type strains.

**Table 1.**

| Strain | Age | Sample | Mean ± SD | | | |
| --- | --- | --- | --- | --- | --- | --- |
|  |  |  | Telomere | Reactive oxygen species (ROS) | Mitochondria viability | Lipid peroxidation |
| | | | aTL<br>(Mb/diploid genome) | DCFH<br>(nM) | MTT<br>(μM) | MDA<br>(μM) |
| Wt | 5 | Hippocampus | 172.1 ± 132.9 | 100.8 ± 67.6 | 0.08 ± 0.03 | 0.50 ± 0.17 |
| | | Whole blood cells | 131.3 ± 22.7 | 635.7 ± 158.9 | ----- | 0.07 ± 0.05 |
| | 9 | Hippocampus | 78.8 ± 28.7 | 215.2 ± 40.3 | 0.03 ± 0.01 | 0.94 ± 0.28 |
| | | Whole blood cells | 133.2 ± 124.2 | 1295.0 ± 252.5 | ----- | 0.09 ± 0.06 |
| | 13 | Hippocampus | 134.0 ± 85.6 | ----- | ----- | ----- |
| | | Whole blood cells | 96.7 ± 60.5 | ----- | ----- | ----- |
| Tg | 5 | Hippocampus | 106.9 ± 64.4 | 444.5 ± 197.3 | 0.10 ± 0.01 | 0.35 ± 0.14 |
| | | Whole blood cells | 149.0 ± 20.9 | 1777.2 ± 339.5 | ----- | 0.10 ± 0.04 |
| | 9 | Hippocampus | 104.8 ± 62.3 | 1442.3 ± 1395.5 | 0.03 ± 0.01 | 1.25 ± 0.38 |
| | | Whole blood cells | 153.4 ± 64.7 | 3796.1 ± 1061.9 | ----- | 2.81 ± 1.49 |
| | 13 | Hippocampus | 52.2 ± 49.5 | ----- | ----- | ----- |
| | | Whole blood cells | 15.4 ± 9.3 | ----- | ----- | ----- |
| Mean ± SD | | | | | | |

aTL, absolute telomere length; Mb, megabase.

### Telomere length

Telomere length in whole blood cells of the transgenic strain showed significant differences between age groups (KW; H(2) = 14.01, p < 0.001), the 13-month-old group was significantly different from the other two groups (Dunn’s post hoc; p = 0.014, p = 0.002). No significant differences were found between age groups from the wild type strain (KW; H(2) = 0.70 p= 0.70). When compared between mice strains, significant differences were found between 13-month-old transgenic and wild type mice (Dunn’s post hoc; p = 0.016) as can be seen in Figure 1a.

**Figure 1.**
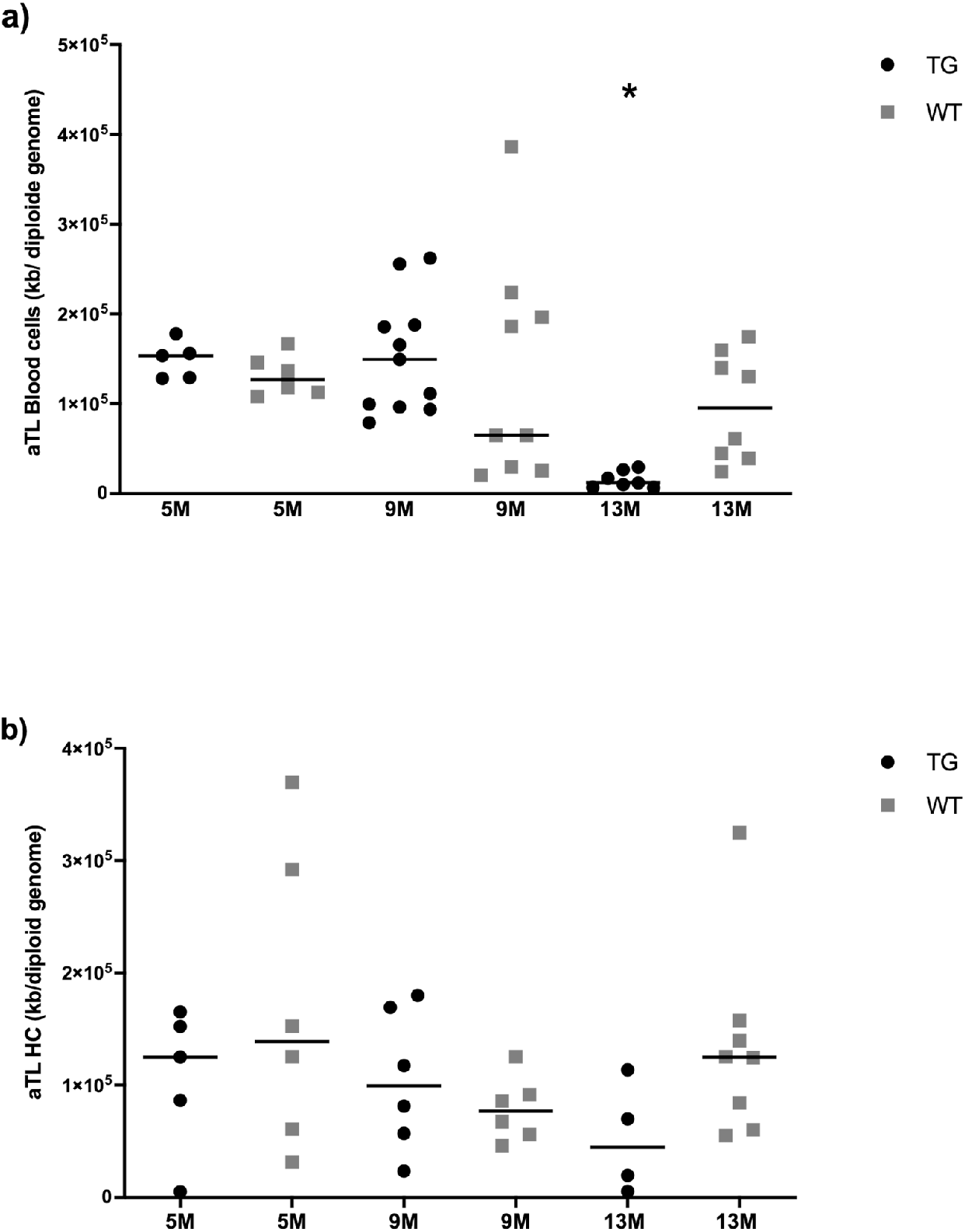
Absolute telomere length measured in transgenic (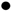black) and wild type (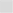grey) mice of 5, 9 and 13 months of age. a) Blood cells telomere length decreases with age and is significant between 5 and 13 and 9 and 13 months of age in transgenic mice, but no significant changes were found for wild type. Significant differences between strains of 13 months of age were found. b) Hippocampus telomere length is not affected by strain or age group. * p <0.05

Telomere length was also measured in the hippocampus, one of the first structures affected by AD. We observed a telomere length decreasing tendency for HC of transgenic mice; nevertheless, we did not find significant differences between age groups within strains (KW; H(2) = 2.45 p = 0.31 for transgenics and KW; H(2) = 2.02 p = 0.37 for wild type) or between wild type and transgenic mice (KW; H(6) = 5.58 p = 0.34) as can be seen in Figure 1b.

### Oxidative stress

In order to determine whether oxidative stress increased with the progression of AD, we measured ROS (by DCFH) and lipid peroxidation (by MDA) at 5 and 9 months of age. When measured in blood cells, both ROS and lipid peroxidation showed a significant increase between age groups within strains. As shown in Figure 2a, a Kruskal wallis test show differences between groups (KW; H (3) = 18.79 p < 0.001) ROS in blood cells of 9 months old transgenic mice is significantly higher than that of 5 months old mice (Dunn’s post-hoc; p < 0.001). ROS also increased between 5 and 9 months old groups of the wild type strain (Dunn’s post-hoc; p = 0.002). ROS increases with age in both strains, but significantly more in the transgenic mice where the 9 months group also showed differences between strains (Dunn’s post-hoc; p = 0.021).

**Figure 2.**
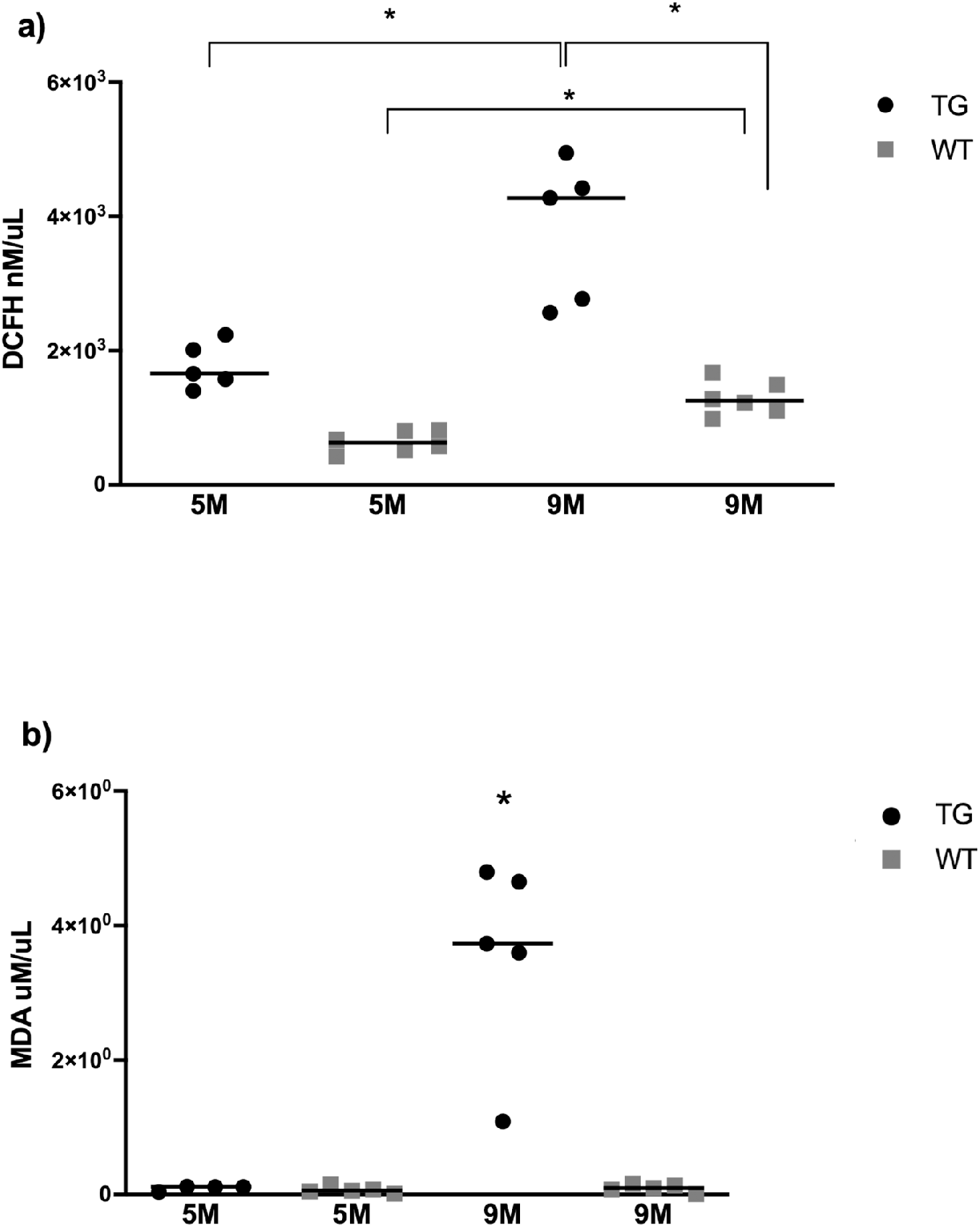
Oxidative stress in blood cells of transgenic (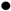black) and wild type (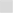grey) mice of 5 and 9 months of age. a) Measurement of ROS by DCFH. Blood cells ROS increases with age in both strains and is statistically significant at 9 months of age between strains. b) Lipid peroxidation measured by MDA. Significant differences were found between 9-month-old transgenic mice and every other group. * p <0.05

Lipid peroxidation, measured by MDA (μM/μL), is shown in figure 2b, where increases can only be seen in the 9 months-old transgenic mice group which is significantly different from every other group (KW; H(3) = 10.99 p = 0.01). No other significant differences in lipid peroxidation were found.

When measured in the hippocampus, ROS increased with age within both strains, a Mann Whitney test showed differences between wild type (Mdn = 83.30, Mdn= 210.3, U = 4 p = 0.026) and transgenic mice in both ages (Mdn = 427.5, Mdn= 905.3, U = 2 p = 0.032) moreover we could observe that increase in ROS was more in transgenic mice at 9 months than wild type mice at the same age (Mdn = 905.3, Mdn= 210.3, U = 0 p = 0.004) as can be seen in Figure 3a. In lipid peroxidation significant increase between 5 and 9 months old mice for both strains, transgenic (Mann Whitney; Mdn = 0.284, Mdn= 1.131, U = 0 p = 0.008) and wild type (Mann Whitney; Mdn = 0.495, Mdn= 0.874, U = 3 p = 0.015) however no differences were observed between strains (Figure 3b). Similarly, mitochondrial viability (measured by MTT) decreased with age in both strains, transgenic mice (Mann Whitney; Mdn = 0.098, Mdn= 0.027, U = 0 p = 0.008) and wild type mice (Mann Whitney; Mdn = 0.076, Mdn= 0.028, U = 0 p = 0.002) but showed no differences between strains as shown in Figure 3c.

**Figure 3.**
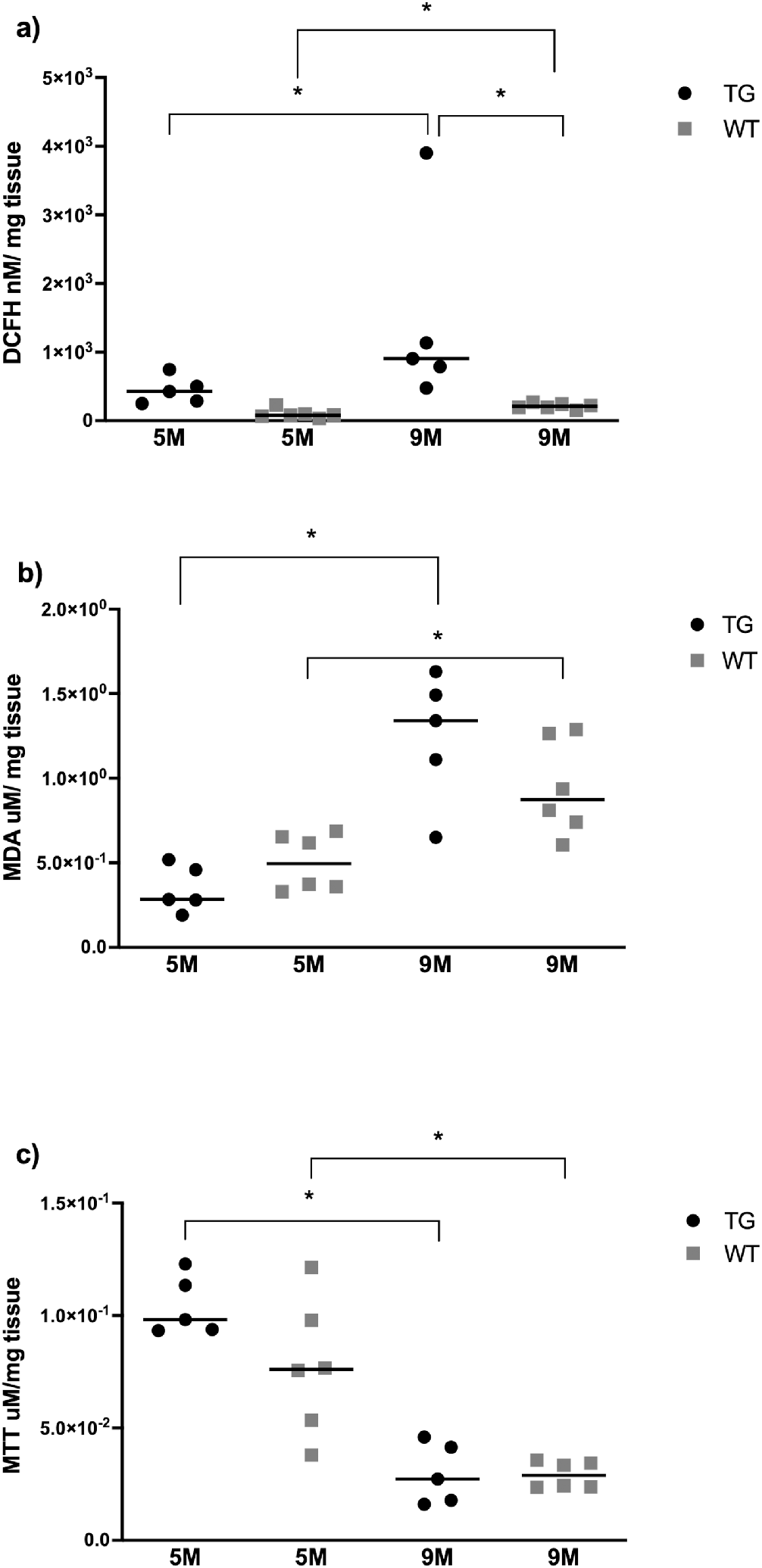
Oxidative stress in hippocampus homogenates of transgenic (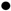black) and wild type (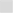grey) mice of 5 and 9 months of age. a) Measurement of ROS by DCFH. ROS increases with age in both strains and is statistically significant at 5 and 9 months of age between strains. b) Lipid peroxidation measured by MDA. Significant differences were found between 5 and 9-month groups of both strains but not between strains. c) Mitochondrial viability measured by MTT. Significant differences were found between 5 and 9-month groups of both strains but not between strains. * p <0.05

Finally, a correlation analysis showed a low/moderate positive correlation for 5 month-old wild type mice between telomere length and ROS for the hippocampus (HC rs=0.3) and whole blood cells (WB rs=0.5) as well as for MDA (HC rs=0.3 and WB rs=0.7), while a low negative correlation was found for MTT in the hippocampus (HC rs= −0.5). Similar correlations were seen for the 9 month-old wild type mice with the exception of a low negative correlation between HC telomere length and MDA (rs= −0.1) and a high positive correlation between WB telomere length and MDA (rs=0.9, p=0.04) as can be seen in Figure 4.

**Figure 4.**
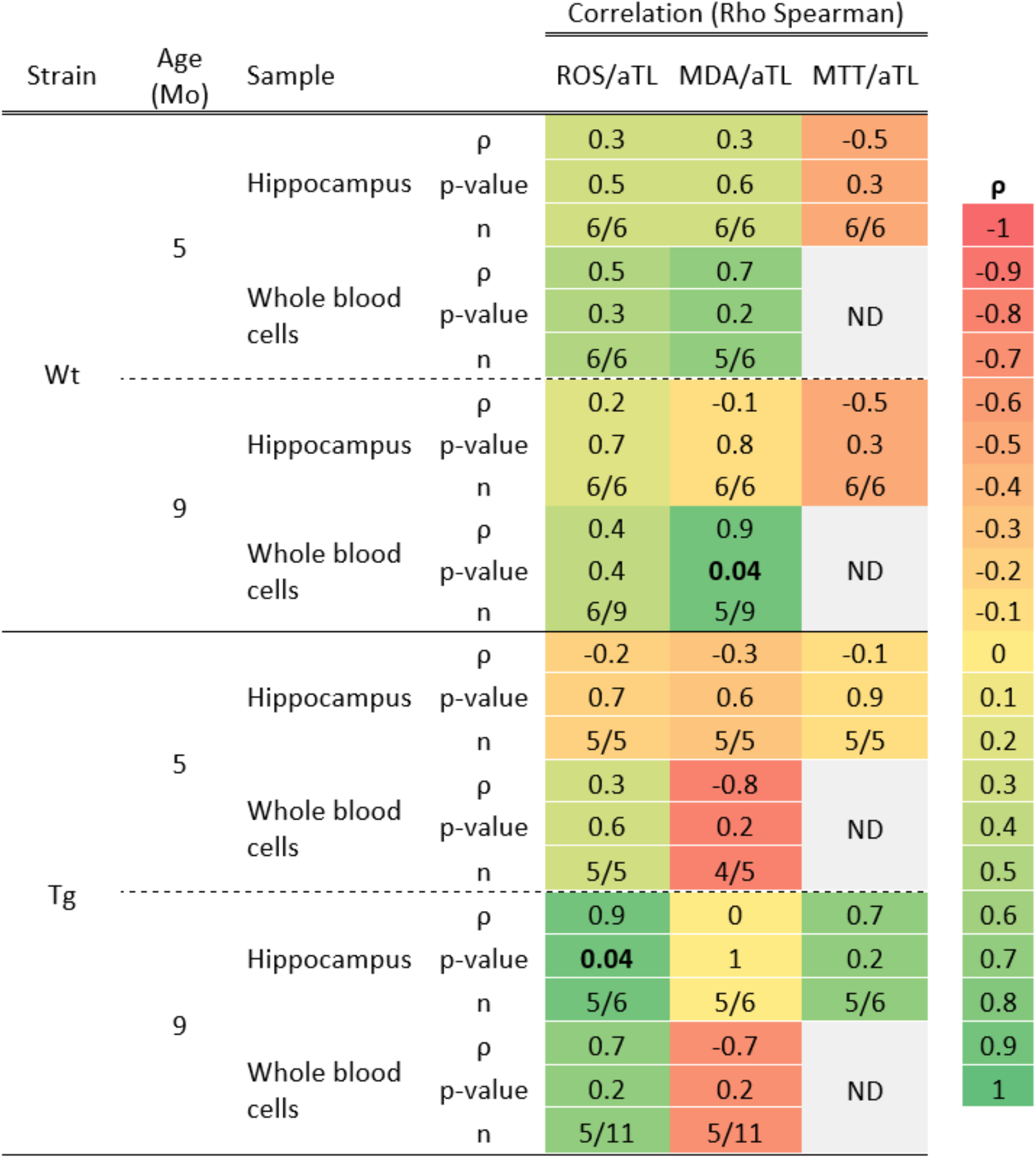
Correlation between telomere length and oxidative stress by strain, age and tissue sample. The combined table/heatmap shows the correlation between oxidative stress (ROS, MDA, and MTT) and telomere length in the hippocampus and whole blood cells from wild-type and transgenic mice at 5 and 9 months of age in a color scale from −1 to +1. ρ, Spearman rho; aTL, absolute telomere length, Wt, wild-type; Tg, transgenic; Mo, months

For the transgenic strain, the hippocampus from 5 month-old mice, showed low negative correlations between telomere length and ROS, MTT, and MDA (HC rs= −0.2, −0.1, and −0.3, respectively) and high negative correlation for whole blood cells and MDA (rs= −0.8). Finally, 9 month-old mice showed moderate and high positive correlations between HC telomere length and ROS (rs= 0.9, p=0.04) and MTT (rs= 0.7), respectively, while positive and negative moderate correlations for whole blood cells with ROS (rs= 0.7) and MDA (rs= −0.7) respectively as can be seen in Figure 4.

## Discussion

Telomere length and oxidative stress are affected by AD progression in the 3xTg-AD murine model. Shorter telomeres were found in blood cells of older transgenic mice compared to younger transgenic mice and same-age wild type mice. This could be a reflection of a systemic effect due to the progression of AD in our murine model since it is not aging but the disease itself that is causing the changes in telomere length.

On the matter, it has been reported that chronic inflammation promotes telomere attrition by increasing white blood cell replacement (Samani *et al.*, 2001). An increase of glial fibrillary acidic protein immunoreactivity, indicative of astrogliosis, has been reported in the retinal ganglion cell layer in the late-symptomatic stages of 3xTg-AD (Edwards *et al.*, 2014). During disease progression in the 3xTg-AD mouse model, retinal microglia showed a pro-inflammatory phenotype, with less ramified morphology, and neurodegenerative associated markers (Grimaldi *et al.*, 2018). Hence, inflammation could promote white blood cell replacement which would reflect in telomere shortening for this murine model.

Additionally, it has been shown that elevated levels of Aβ 1-40 and Aβ 1-42 are associated with increased oxidation products in peripheral blood cells from AD patients (Coppedè & Migliore, 2015). On the matter, plasma levels of Aβ 1-40 and Aβ 1-42 of 3xTg-AD mice have been found to increase progressively from 5 to 9 months of age (age at which this model does not yet show accumulation of Aβ in brain tissue), observing a significant decrease at 12 months (Cho *et al.*, 2016).

Hence, the elevation of oxidative stress that we see with age, and that is more evident and prominent in the transgenic model, could be due to the presence of Aβ in blood, which appears sooner than in brain tissue. However, at the time we cannot statistically prove that the elevated oxidative stress causes telomere attrition, even when it seems a likely scenario.

We found increased oxidative stress, measured by ROS and lipid peroxidation, in blood cells of older compared to younger transgenic mice; however, we found no statistical association between the increase in oxidative stress and telomere length in our model. For that purpose, 8OHdG may have been a better biomarker to associate with telomere length since it is directly related to DNA damage due to oxidative stress. Accordingly, a previous paper reported that 8OHdG is elevated in lymphocytes of AD patients compared to a control group (Mecocci *et al.*, 1994). Unfortunately, this measurement was impossible to obtain at the time, and the latter study was made in humans.

Regarding our findings on telomere length in the hippocampus (HC), we only found a tendency, but no significant differences between the transgenic 13 months old mice and the younger groups. On that matter, Franco and co-workers found no changes in telomere length of the subcortical and granular areas as well as the HC between transgenic mice for APP and their controls. They concluded that the accumulation of amyloid-beta has no relation to telomere shortening (Franco *et al.*, 2006). Moreover, Lukens et al. found no changes in telomere length in the cerebellum of patients with AD against controls despite a significant difference in leukocyte telomere length (Lukens *et al.*, 2009a). However, Thomas et al., 2008 found a significant increase in telomere length in the hippocampus of brain tissue of AD patients compared to controls as well as a significantly shorter telomere of the younger AD compared to older AD patients (Thomas *et al.*, 2008). Finally, Cardillo et al., 2018 found no differences in the hippocampal telomere length of 3xTg-AD 11-month-old mice compared to WT controls (Cardillo *et al.*, 2018)

Changes in telomere length in brain tissue are tricky to interpret since most neurons do not replicate and specifically, the hippocampus shows reduced neurogenesis in AD (Dhaliwal *et al.*, 2018). A recent stereological study in eleven-month-old 3xTg-AD mice indicated that, in spite of the occurrence of cerebral atrophy and reduced hippocampal volume, there was preservation of the total number of CA1 pyramidal neurons, suggesting a self-preservation mechanism that may be reflected in enhanced telomere maintenance (Manaye *et al.*, 2013; Schaeffer *et al.*, 2017). Therefore, if shortening occurs, it is more likely to be due to environmental factors or to increased replication of glial cells (Thomas *et al.*, 2008; Cattan *et al.*, 2008), a distinction we did not make at the moment.

Here we found that the progression of the disease does not affect telomere length in brain tissue as it does on blood cells. On the matter, it has been reported that telomere length in skeletal muscle (a minimally replicating somatic tissue), and leukocytes (a highly proliferative hematopoietic system) shorten with age, but only leukocytes show statistically significant attrition with time (Chahine *et al.*, 2019). Telomeres in blood cells may shorten at a greater rate than other tissues because of their high turnover rate (Nakagawa *et al.*, 2004). Hence, inter-species, interindividual, and intertissue differences are accounted for mainly by factors such as different rates of cell replication, levels of telomerase activity, and levels of oxidative stress (Aviv, 2002).

On that matter, it is worthy to note, that here we did not take into account the effects of interstitial telomeric sequences (ITSs). The dynamics for telomere estimation are complex because telomeric repeats exist both within true telomeres at the ends of chromosomes and as ITSs in the interior of chromosomes. Quantitative PCR (Q-PCR) detects both true telomere and ITSs (Nakagawa *et al.*, 2004; Foote *et al.*, 2013). ITSs are likely to affect telomere estimates in any species that has substantial ITSs relative to true telomeres. This is problematic since it could increase the variance within a group of samples, as we can see in our measurements. Both of these effects reduce statistical power to detect significant differences in telomere length between groups (Foote *et al.*, 2013).

Regarding aging and telomere shortening, a negative correlation between age and telomere length (Lukens *et al.*, 2009) has been reported in human tissue and peripheral blood cells, a feature we did not find in any of our mice, wild type or transgenic. Accordingly, a comparative study in mice established no correlation between telomere length and age in these animals; the replicative aging mechanism controlling the number of cell divisions where telomere length is shortened, as has been reported in humans, appears to be different in mice (Gomes *et al.*, 2011). Murine telomeres do not serve as a mitotic clock for replicative aging, as primary cells constitutively express telomerase, in contrast to humans, in whom telomeres play a part in replicative senescence and telomerase expression is repressed (Calado & Dumitriu, 2013; Reichert & Stier, 2017). Hence, the data obtained here must be taken with caution when compared to humans.

Taking it all into account, human and mice studies have both described no changes in telomere length in brain tissue samples but a shortening in telomere in whole blood cells of AD patients. Although oxidative damage can cause telomere shortening through double-stranded breaks to DNA, most telomere loss due to oxidative stress occurs during DNA replication (Calado & Dumitriu, 2013; Reichert & Stier, 2017). Oxidation of biomolecules in the context of AD is mainly related to neuronal membrane biomolecules and to a disruption of membrane integrity (Cheignon *et al.*, 2018), which is in accordance to our findings of a general negative correlation found between lipid peroxidation (measured by MDA) and telomere length for both brain tissue and whole blood cells of transgenic mice. This correlation is mainly evident and statistically significant for whole blood cells of transgenic 5 month-old mice.

Our results show that telomere attrition is due to the progression of the disease, supporting the newest discovery of AD being a disease that does not only affect the central nervous system but has a systemic effect.

Aging, as well as AD, has an effect on oxidative stress (Panossian, 2003; de Souza-Pinto *et al.*, 2008) which we were able to confirm here. Both mice strains had increased markers of oxidative stress for blood and tissue samples when compared to a younger group, but transgenic mice showed even higher values suggesting an additive effect of age and Alzheimer’s Disease. However, no statistical correlation between oxidative stress increase and telomere length was found. Independently of age, our results point to a distinct, not ROS-related, and yet to be determined, the global mechanism by which the Alzheimer phenotype promotes cell proliferation and consequently telomere shortening.

## Conclusion

Higher oxidative stress and shorter telomere length in peripheral blood were observed with the progression of the disease but no statistical correlation between them was found. No changes in telomere length but an increase in oxidative stress was found in the hippocampus of both strains. The presence of changes in peripheral blood cells of this mouse model suggests that AD affects the individual systemically due to its progression.

## Acknowledgments

This work was supported by the Fondo Sectorial de Investigación para la Educación del Consejo Nacional de Ciencia y Tecnología (CONACyT) with the project number CONACYT 179594.

We would like to thank Ariadna Galvan Flores for her help with the design and elaboration of the graphical abstract and figures.

This study was performed in partial fulfilment of the requirements for the degree of KMG (MSc) and AIH (PhD) who thank the Posgrado en Ciencias Biológicas, Biología Experimental, and acknowledge the scholarships provided by CONACYT (294248 and 298982 respectively) and KMG for IMSS (99096757).

## Abbreviations

Aβ: amyloid beta
aTL: absolute telomere length
AD: Alzheimer’s disease
DNA: deoxyribonucleic acid
DCFH: 2-7-dichlorofluorescein
DCF: oxidized compound of DCFH
HC: hippocampus
ITSs: interstitial telomeric sequences
MDA: malondialdehyde
mo: months
MTT: 3-(4,5-dimethylthiazol-2-yl)bromide-2,5-diphenyltetrazole
qPCR: quantitative polymerase chain reaction
ROS: reactive oxygen species

## Notes

### Competing Interest Statement

The authors have declared no competing interest.

